# Systematic comparison of incomplete-supervision approaches for biomedical imaging classification

**DOI:** 10.1101/2020.12.07.414235

**Authors:** Sayedali Shetab Boushehri, Ahmad Bin Qasim, Dominik Waibel, Fabian Schmich, Carsten Marr

## Abstract

Deep learning based classification of biomedical images requires manual annotation by experts, which is time-consuming and expensive. Incomplete-supervision approaches including active learning, pre-training and semi-supervised learning address this issue and aim to increase classification performance with a limited number of annotated images. Up to now, these approaches have been mostly benchmarked on natural image datasets, where image complexity and class balance typically differ considerably from biomedical classification tasks. In addition, it is not clear how to combine them to improve classification performance on biomedical image data.

We thus performed an extensive grid search combining seven active learning algorithms, three pre-training methods and two training strategies as well as respective baselines (random sampling, random initialization, and supervised learning). For four biomedical datasets, we started training with 1% of labeled data and increased it by 5% iteratively, using 4-fold cross-validation in each cycle. We found that the contribution of pre-training and semi-supervised learning can reach up to 25% macro F1-score in each cycle. In contrast, the state-of-the-art active learning algorithms contribute less than 5% to macro F1-score in each cycle. Based on performance, implementation ease and computation requirements, we recommend the combination of BADGE active learning, ImageNet-weights pre-training, and pseudo-labeling as training strategy, which reached over 90% of fully supervised results with only 25% of annotated data for three out of four datasets.

We believe that our study is an important step towards annotation and resource efficient model training for biomedical classification challenges.

## Introduction

Recent successes of deep learning methods rely on large amounts of well annotated training data^1^. However, for biomedical images annotations are often scarce as they crucially depend on the availability of trained experts, whose time is expensive and limited. Therefore many biomedical imaging classifications can be categorized as incomplete-supervision approaches, where the labeled data is limited and unlabeled data is abundant^2^. While encountering an incomplete-supervision problem, two questions typically arise: How much annotation is needed to train a decent model? And which computational methods can be applied to arrive there efficiently?

Active learning algorithms address the issue by finding the most informative instances for annotation^3–5^ and have been benchmarked on natural image datasets^6–13^. Pre-training methods such as transfer learning and self-supervised learning have shown a great potential for improving the network performance on classification tasks with only a small number of labeled images^14–17^. During transfer learning, a neural network uses the representation from another model, ideally trained on a similar dataset, while in self-supervised learning, a representation without any labels is learned^18^. A common transfer learning approach, used also in many biomedical applications, is to initialize a model with pre-trained ImageNet weights^19,20^. Semi-supervised learning leverages unlabeled data in addition to labeled data to increase the performance as well as the stability of predictions^21,22^. This is particularly appealing for biomedical imaging, where high-throughput technologies^23^ generate large quantities of unlabeled data.

Biomedical image datasets differ from natural images in a couple of important characteristics: They are often imbalanced, typically less diverse in terms of shapes and color range, and classes are often distinguished by only small feature variations, e.g. in texture and size^24,25^. While there is an increasing number of publications covering these three approaches separately, it is not clear which combination of the aforementioned approaches yields the best performance in practice. Moreover, the approaches’ efficiency with respect to each other has not been analyzed.

In this paper, we perform an extensive grid-search in incomplete-supervision approaches including seven active learning algorithms plus random sampling as the baseline, three pre-training methods plus random initialization as the baseline, and training strategies including two semi-supervised learning methods and supervised learning as the baseline, on four exemplary biomedical imaging datasets. First, we compare which combination leads to the best results on each dataset. For each dataset, we then analyze the contribution of the best active learning algorithm, best pre-training method and best training strategy. Finally, we recommend a combination of approaches for dealing with similar biomedical classification tasks.

## Results

### Biomedical image datasets

To evaluate the efficiency and performance of active learning algorithms, pre-training methods, and training strategies, we have selected four exemplary, publicly available and fully annotated datasets from the biomedical imaging field. These datasets show strong class imbalance, little color variance, strong similarity between classes, and cover different applications (Figure 1):

**Figure 1.**
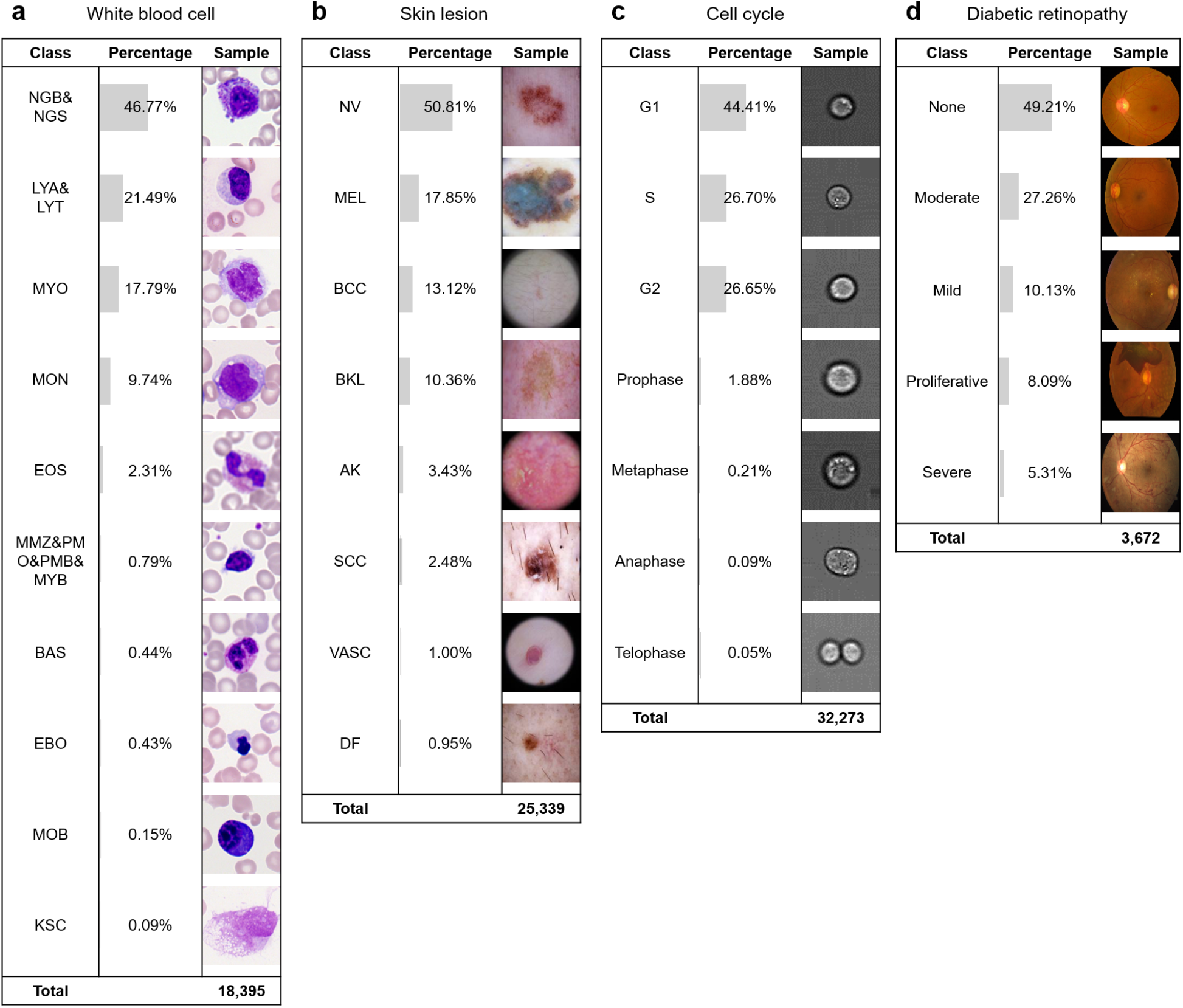
Biomedical image datasets exhibit strong class imbalance, little color variance and high similarity among classes. (**a**) The white blood cell dataset comprises 18,395 images (128×128×3 pixel) of human leukocytes from blood smears of 100 patients diagnosed with Acute Myeloid Leukemia and 100 individuals who show no symptoms of the disease^24,26,33^ with ten expert labeled classes. (**b**) The skin lesion dataset contains 25,339 dermoscopy images (128×128×3 pixel) from eight skin cancer classes^27–29^. (**c**) The cell cycle dataset comprises 32,273 images (64×64×3 pixel) of Jurkat cells in seven different cell cycle stages created by imaging flow cytometry^30^. For better visualization, only the bright-field channel is shown. (**d**) The diabetic retinopathy dataset consists of 3,672 color fundus retinal photography images (128×128×3 pixel) classified into five stages of diabetic retinopathy^31^. All datasets are publicly available (see Methods).

- The white blood cell dataset comprises 18,395 images (128×128×3 pixel) of human leukocytes from blood smears of 100 patients diagnosed with acute myeloid leukemia and 100 individuals who show no symptoms of the disease^24,26^, with 15 expert labeled classes. To ensure a meaningful test set, we have merged neutrophils (segmented and band), lymphocytes (typical and atypical) and immature leukocytes (myeloblasts, promyelocytes, promyelocytes-bilobed, and myelocytes) based on the class definitions^24^ (Figure 1a).
- The skin lesion dataset contains 25,339 dermoscopy images (128×128×3 pixel) from eight skin cancer classes^27–29^. The dataset has been used in the ISIC 2018 challenge as an effort to improve melanoma diagnosis (Figure 1b).
- The cell cycle dataset comprises 32,273 images (64×64×3 pixel) of Jurkat cells in seven different cell cycle stages captured by imaging flow cytometry^30^. The four minority classes cover only 2.4% of the data. In addition, there is a great amount of similarity among the classes (Figure 1c).
- The diabetic retinopathy dataset consists of 3,672 high-resolution color fundus retinal photography images^31^, classified into 5 stages of diabetic retinopathy. For computational reasons, we have reduced the size of the images from 2095×2095×3 to 128×128×3 pixels. The dataset has been used in the APTOS 2019 Blindness Detection challenge on kaggle^32^ (Figure 1d).

### Experiments

From each dataset, we randomly selected 1% of the data as our initial annotated set and trained a ResNet18 (see Methods). Then in each cycle we added 5% of annotated data as suggested by one of the seven active learning algorithms (Figure 2) or randomly sampled 5% as baseline. This process was repeated eight times which led to adding 40% (and using in total 41%) of annotated data. We combined active learning with three different pre-training methods and random initialization as baseline, and two different training strategies with supervised learning as baseline (see Methods for a detailed description of all methods). This resulted in 13,824 independent experiments (see Figure 2). We performed a 4-fold cross-validation in each cycle and calculated macro F1-score, accuracy, precision, and recall. The macro F1-score, defined as the average F1-score over all classes thus accounting for the imbalanced nature of the datasets (see Figure 1), was used as our main metric of comparison, assuming that the correct prediction of small classes is of biological or diagnostic importance. To quantitatively compare different combinations, we looked at the average macro F1-score across all cycles. Every combination is reported in the form of “active learning algorithm + pre-training method + training strategy”.

**Figure 2:**
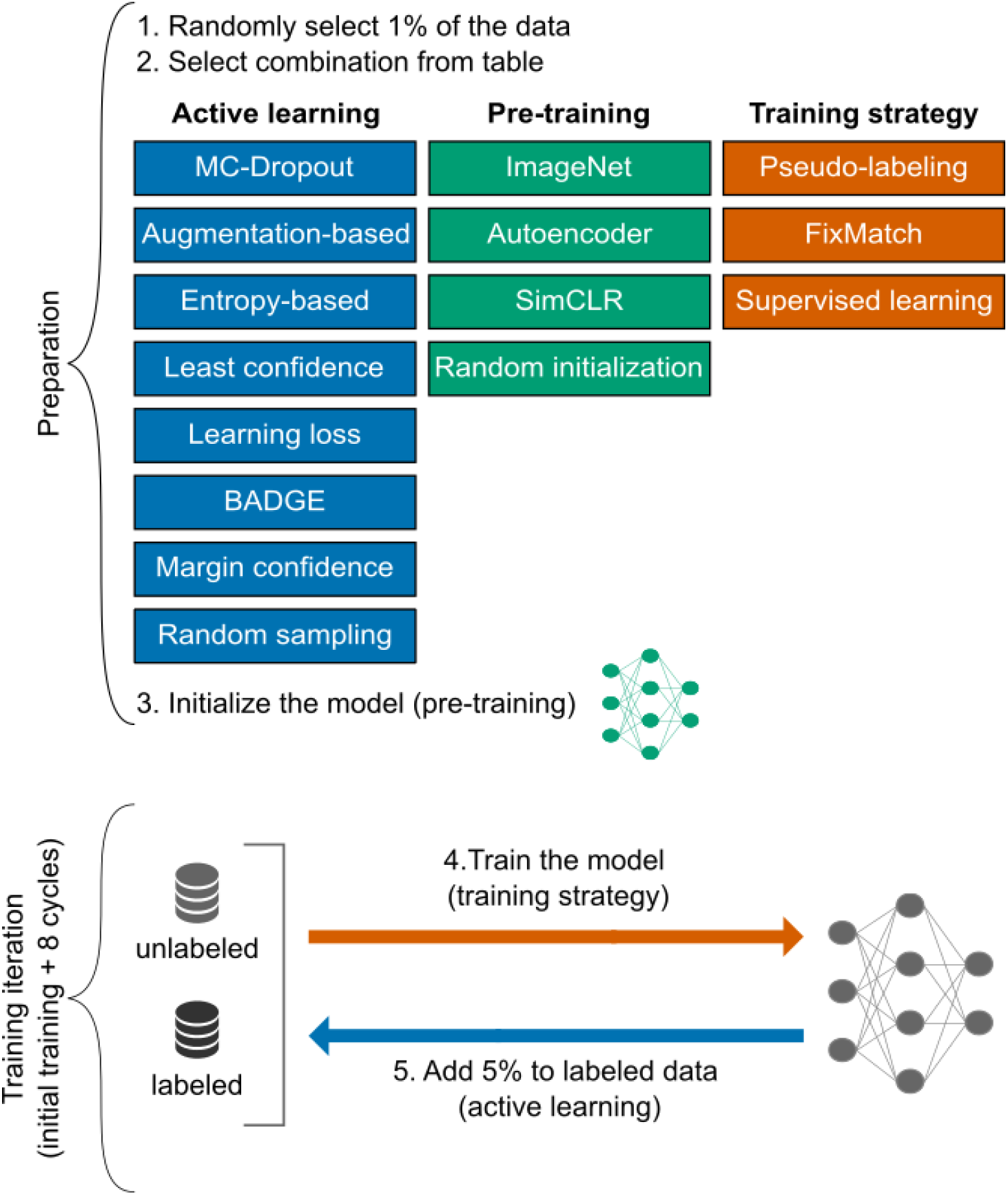
We systematically compared combinations of 7 active learning algorithms, 3 pre-training methods and 2 training strategies on 4 biomedical imaging datasets. Specifically, we ran 4×8×4×3×4×9 = 13,824 independent experiments (4 datasets, 7 active learning algorithms + 1 baseline, 3 pre-training methods + 1 baseline, 2 training strategies + 1 baseline, 4-fold cross-validation and 1 initial step + 8 active learning cycles) to identify the best combination out of 96 possible combinations.

The best performing combination on the white blood cell dataset was learning loss + SimCLR + pseudo-labeling (Figure 3a), which achieved an average macro F1-score of 0.71±0.07 (mean±standard deviation on n=8 cycles). This was followed by entropy-based + SimCLR + pseudo-labeling as well as MC-dropout + SimCLR + pseudo-labeling (0.71±0.08), learning loss + ImageNet + pseudo-labeling (0.71±0.09) and BADGE + ImageNet + pseudo-labeling (0.70±0.09). The best performing combinations reached 94% of the macro F1-score of a model trained in a fully supervised manner on the whole dataset (macro F1-score 0.83±0.02).

**Figure 3.**
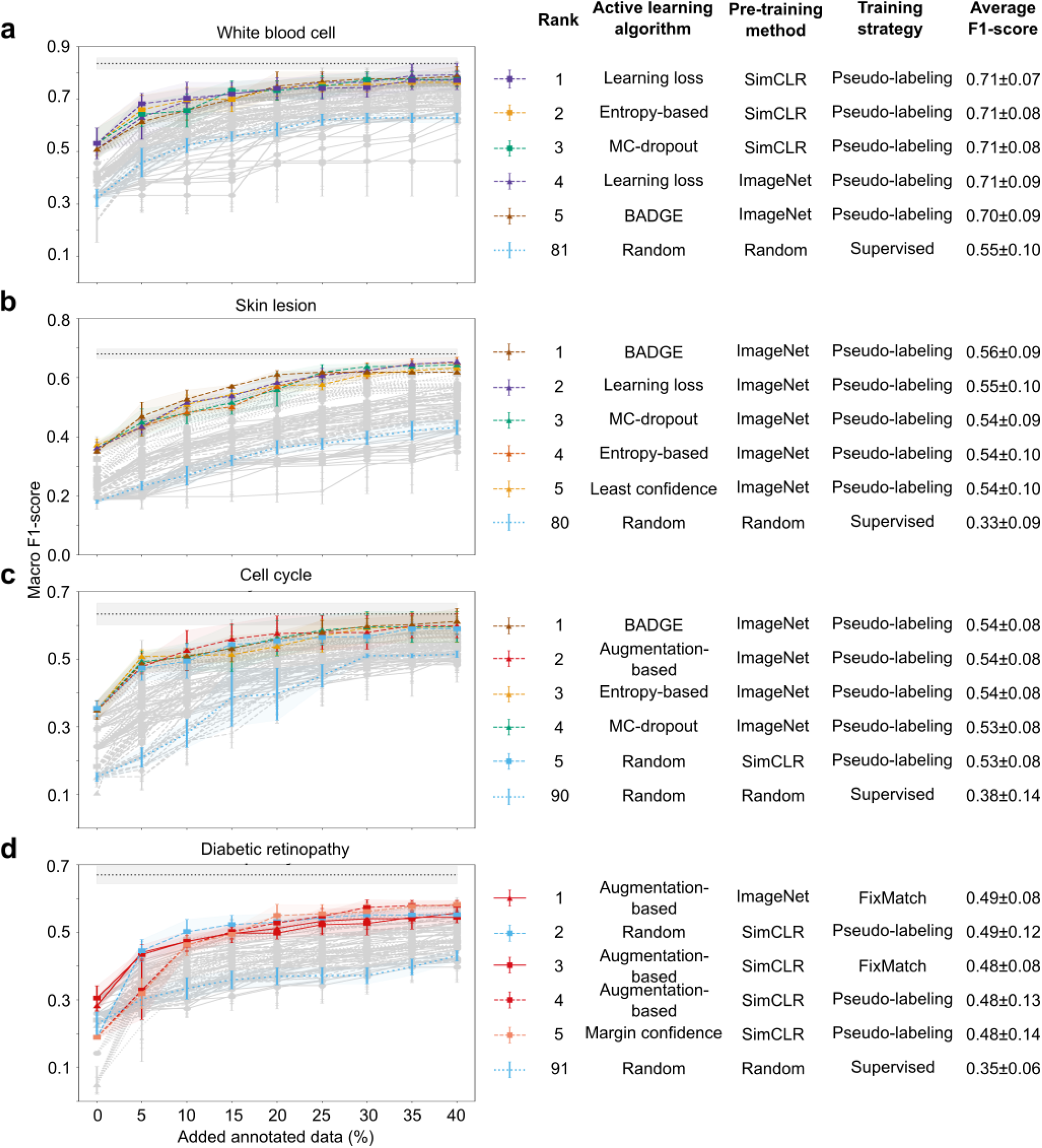
ImageNet and SimCLR as pre-training methods, and pseudo-labeling as the training strategy dominate the best performing combinations, while no particular active learning algorithm prevails. In each panel (a-d) the upper bound of performance is fully supervised learning (black dotted line). (**a**) White blood cell dataset: Pseudo-labeling is the best performing training strategy for this dataset, achieving 94% of the fully supervised results with learning loss and SimCLR. (**b**) Skin lesion dataset: pseudo-labeling and ImageNet pre-training are consistently part of the best combinations, reaching 91% of fully supervised results with BADGE active learning. (**c**) Skin lesion dataset: pseudo-labeling is the top performing training strategy, reaching at least 96% of fully supervised results with augmentation-based active learning and ImageNet pre-training. (**d**) Diabetic retinopathy dataset: random sampling + SimCLR + pseudo-labeling reaches performance similar or better than any other combination, reaching 83% of the fully supervised results.

On the skin lesion dataset BADGE + ImageNet + pseudo-labeling achieved the highest average macro F1-score of 0.56±0.09 (Figure 3b). It was followed by learning loss + ImageNet + pseudo-labeling at 0.55±0.10, MC-dropout + ImageNet + pseudo-labeling at 0.54±0.09, entropy-based + ImageNet + pseudo-labeling as well as least confidence + ImageNet + pseudo-labeling at 0.54±0.10. Notably, the best performing combination reached 91% of the macro F1-score of a model trained in a fully supervised manner on the whole dataset (macro F1-score of 0.68±0.02).

The best performing combinations on the cell cycle dataset were BADGE + ImageNet + pseudo-labeling, augmentation-based + ImageNet + pseudo-labeling, and entropy-based + ImageNet + pseudo-labeling at 0.54±0.08 (Figure 3c). It was followed by MC-dropout + ImageNet + pseudo-labeling as well as random sampling + SimCLR + pseudo-labeling at 0.53±0.08. The best performing combinations reached 96% of the macro F1-score of the same network trained in a fully supervised manner on the whole dataset (macro F1-score of 0.63±0.03).

On the diabetic retinopathy dataset augmentation-based + ImageNet + FixMatch reached the highest macro F1-score at 0.49±0.08, followed by random sampling + SimCLR + pseudo-labeling at 0.49±0.12, augmentation-based + SimCLR + FixMatch at 0.48±0.08, augmentation-based + SimCLR + pseudo-labeling at 0.48±0.13 and margin confidence + SimCLR + pseudo-labeling at 0.48±0.14 (Figure 3d). The best performing combinations reached 83% of the macro F1-score of the same network trained in a fully supervised manner on the whole dataset (macro F1-score of 0.67±0.03).

### Ablation study

In the previous section we made two noteworthy observations. First, no active learning algorithm showed up in the best combinations consistently. Second, ImageNet and SimCLR pre-training as well as pseudo-labeling were always in the top combinations. To better understand each approach’s contribution to the performance, we selected the top combination for each dataset (see Figure 3) and conducted a systematic ablation study. We define the contribution to performance of each incomplete-supervision approach by calculating the difference in F1-score if that approach was substituted with its baseline: active learning algorithms were substituted with random sampling, pre-training methods with random initialization and training strategies with supervised learning (Figure 4).

**Figure 4.**
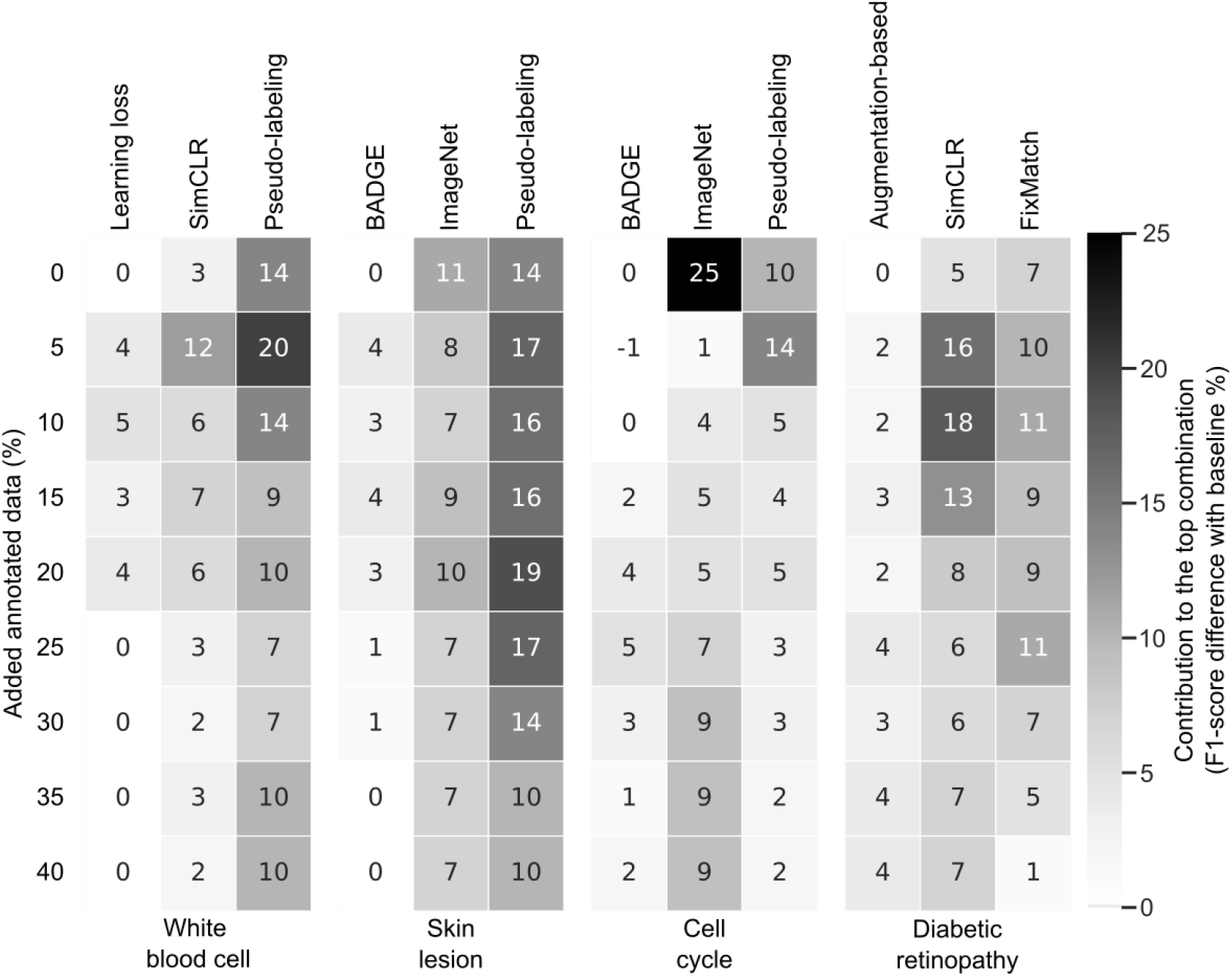
Semi-supervised learning and pre-training contribute stronger to the top performing combination in comparison to active learning. For every dataset, the top combination of active learning algorithm, pre-training method and training strategy based on the results of the grid-search (see Results and Figure 3) is depicted. The contribution to performance of each approach is calculated by substituting it with its baseline and subtracting the obtained macro F1-score from the original (see Methods for details).

Our analysis revealed that although combining active learning, pre-training and semi-supervised learning exhibited to be effective in all datasets, their contributions to performance differ. For every top combination, the contribution of the pre-training and semi-supervised learning is considerably higher than active learning. For example, after adding only 5% annotated data for the white blood cell dataset, learning loss as active learning contributes 4% to the performance, while SimCLR pre-training and pseudo-labeling as training strategy contribute 12% and 20% respectively (2nd row in first matrix in Figure 4). The same kind of observation can be deduced for the other three datasets. Interestingly, the highest contributions to macro F1-score are always achieved in the first 5 cycles.

## Discussion

We have investigated the effect of combining different incomplete-supervision approaches on four biomedical image classification datasets with an extensive grid-search over seven active learning algorithms, three pre-training methods and two training strategies as well as their baselines. For three out of four datasets, the top combinations reached more than 90% in macro F1-score of the fully supervised approach (see Figure 3 and 5b), with only 26% of the data being labeled (1% randomly selected and 5% added in 5 active learning cycles). Notably, this was not the case for the diabetic retinopathy dataset, where the top combination still lacked 12% from the fully supervised results with using even 41% of the labeled data. One reason might be image resolution: For computational reasons, we had to reduce height and width of the images from 2095×2095 to 128×128 pixels, which might have contributed to misclassifications between the ‘proliferate’ and ‘severe’ classes (data not shown).

**Figure 5.**
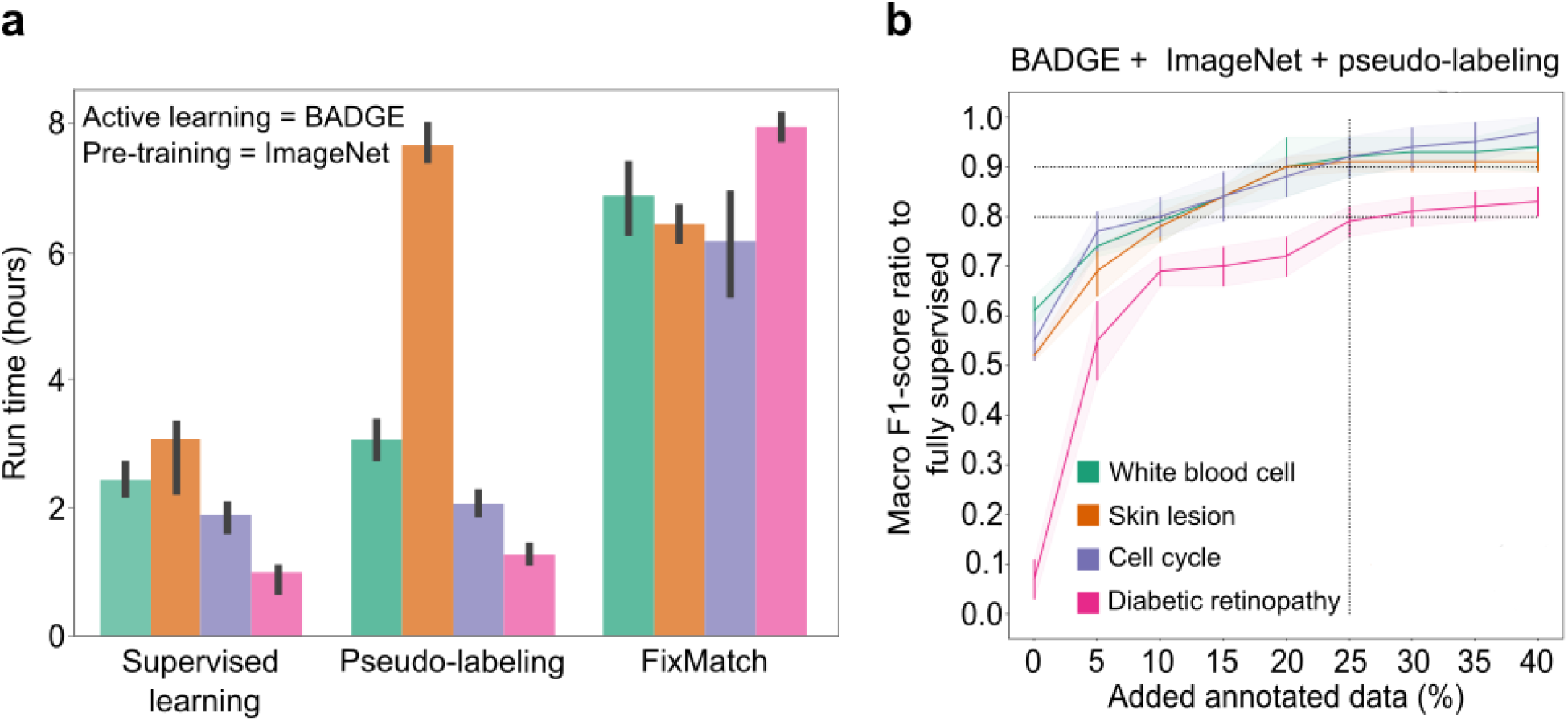
The annotation and resource efficient combination, BADGE + ImageNet + pseudo-labeling, reaches >90% of the fully supervised result in three out of four biomedical datasets by using only 25% of annotated data. (**a**) BADGE + ImageNet + pseudo-labeling takes only slightly longer than supervised learning on three out of four datasets, while BADGE + ImageNet + FixMatch takes at least three times more on every dataset. (**b**) By using only 25% of annotated data, the recommended combination, BADGE + ImageNet + pseudo-labeling, reaches >80% of the fully supervised result for all the datasets and >90% in three out of four datasets.

Our grid-search showed that no single best active learning algorithm outperforms the rest consistently. Even though they perform better than random sampling, the results of using learning loss, augmentation-based, BADGE and MC-dropout are highly dataset dependent. In terms of implementation, all of the methods were straightforward except learning loss, as it brought changes in the architecture, loss function and implementation.

Regarding pre-training methods, ImageNet and SimCLR led consistently to top results, while autoencoder pre-training did not prove to be effective. After close inspection of all combinations (Figure 2 only shows the top-5 combination), we observed that SimCLR showed to be more effective than ImageNet in combination with supervised learning. This observation is in alignment with recent papers that show that SimCLR or other self-supervised methods outperform ImageNet on biomedical applications^34,35,36^. However, our analysis showed that ImageNet and SimCLR pre-training performed comparatively similar when being combined with a semi-supervised method. This can be explained by the fact that semi-supervised learning strategies already use the unlabeled data in their training process, which makes the use of self-supervised methods redundant. In terms of implementation, SimCLR implementation was straightforward, but needed large batch sizes > 2048 which was cumbersome during the execution.

As expected, semi-supervised learning outperformed supervised learning in all cases, due to the fact that it exploits unlabeled data during training. In particular, pseudo-labeling was the top choice for all datasets, while FixMatch only performed well for the diabetic retinopathy dataset. This could be due to the fact that augmentations in FixMatch are not designed for biomedical images (see Methods). While pseudo-labeling implementation was straightforward, tuning the right hyperparameters for FixMatch was rather difficult. In terms of run-time, pseudo-labeling required slightly more than supervised learning, with the exception of FixMatch, which took at least three times more than supervised learning in every case (see Figure 5a).

### Combination

Our analysis showed that combining active learning algorithms, pre-training methods, and semi-supervised learning strategies lead to superior performance in all cases. However, we found that the state-of-the-art active learning algorithms contribute less than 5% to macro F1-score in each cycle. In contrast, the contribution of pre-training and semi-supervised learning can reach up to 25% macro F1-score. Moreover, we found that the initial cycle plays a major role in reaching higher performance. In almost all cases (17 out of 20), the top 5 combinations were the ones that performed well from the first cycle on (see Figure 3). Considering the fact that semi-supervised learning and pre-training contribute more than active learning as well as the importance of the first cycles, we recommend spending more time and resources on testing different semi-supervised learning strategies and pre-training methods instead of active learning.

As a result of this work, we recommend an annotation and resource efficient strategy for biomedical imaging active learning tasks. We propose that the combination of BADGE active learning, ImageNet initialization pre-training, and pseudo-labeling as training strategy can be considered as a stable choice for dealing with problems where annotated data is limited. For three datasets, this combination reached at least 90% of the fully supervised results by only using 25% of the labeled data (see Figure 5b).

Although our work shows the potential of annotation-efficient learning for four biomedical image classification datasets, the methodology should be tested on more datasets to gain insights into correlations between dataset characteristics and the performance of the applied methods. Due to the computational costs, we used a fixed architecture and a fixed set of parameters. While this choice might not lead to the best fully supervised performance for each dataset (e.g. compared to much bigger architectures or series of ensemble learners used in ISIC 2018 and APTOS 2019), it provides a framework to systematically analyze the combination of incomplete-supervision approaches. Based on the work of Chen et al.^37^, we also suggest testing bigger architectures to figure out if there is a correlation between the architecture size and the performance for biomedical data. Finally, a larger variety of active learning algorithms, self-supervised methods and semi-supervised strategies should be added to this analysis to find an overall optimal strategy.

## Methods

### Active learning algorithms

The performance of a model *f*^⊝^ with parameters ⊝ can be increased by labeling images from the set of unlabeled images *U*, and thus adding pairs of images and corresponding labels (*x*_*i*_, *y*_*i*_) to the set of labeled images *L*. The labeling of unlabeled images is carried out in cycles, in which *s* images *S* ⊆ *U* with |*S*| = *s* are selected for annotation and added to *L*, after the performance of the model converges with the previous labeled set *L*. Active learning algorithms aim on selecting images in *U* for annotation, such that the addition of these images to *L* results in a maximum increase in the evaluation metrics *M*. The main difference between active learning algorithms is how images in the *U* are prioritised for labeling. The algorithms evaluated in this paper are based on uncertainty δ. Uncertainty δ is a scalar value which is attributed to each image in *U*. The *s* images *S* ⊆ *U* with |*S*| = *s* with the highest uncertainty are selected for labeling in each cycle.

### Monte Carlo dropout (MC-dropout)

Dropout is a commonly used technique for model regularization, which randomly ignores a fraction of neurons during training to mitigate the problem of overfitting. It is typically disabled during test time. MC-dropout involves the assessment of uncertainty in neural networks using dropout at test time^38,39^ and thus estimates the uncertainty of the prediction of an image. MC-dropout generates non-deterministic prediction distributions for each image. The variance of this distribution can be used as an approximation for model uncertainty δ^40^. During each active learning cycle, the s images with the highest variance are annotated and added to the labeled set *L*. This has been shown to be an effective selection criterion during active learning^6^.

### Augmentation-based sampling

Let *a* be a function that performs stochastic data augmentation, such as cropping, horizontal flipping, vertical flipping or erasing on a given image. Each unlabeled image *u*_*i*_ ∈ *U* is transformed using *a* and this process is repeated *J* times to obtain the set *U*_*i*_ = {*u*_*1i*_, *u*_*2i*_, *u*_*3i*_…*u*_*Ji*_} with |*U*_*i*_| = *J*. The random transformations are followed by a forward-pass through the model *f*^*⊝*^. This results in *J* predictions *Q*^_*i*_ = {*q*^_*1i*_, *q*^_*2i*_, *q*^_*3i*_…*q*^_*Ji*_}, where *q*^_*i*_ = argmax *P*^⊝^(*y*^_*i*_|*u*_*i*_) is the most probable class according to the model output for each set *U_i_* of perturbed copies of an unlabeled image *u*_*i*_ ∈ *U*. The model uncertainty δ can be estimated by keeping a count of the most frequently predicted class (mode) for each image. The idea behind this approach is that if the model is certain about an image, it should output the same prediction for randomly augmented versions. Thus, the lower the frequency of the mode, the higher the uncertainty δ^41^. During each active learning cycle, the images with the lowest frequency of the most frequently predicted class are annotated and added to the labeled set *L*.

### Entropy-based sampling

Entropy measures the average amount of information or “bits” required for encoding the distribution of a random variable^3^. Here, entropy is used as a criteria for active learning^3^ to select the *s* images *S* ⊆ *U*, whose predicted outcomes (softmax layer) have the highest entropy, assuming that high entropy of predictions mean high model uncertainty δ. By definition, entropy focuses on taking the complete predictive distribution into account^3^.

### Least confidence

Least confidence sampling is the simplest and most common form of uncertainty sampling. The difference between the most confident prediction out of all class predictions (the highest softmax value) and 100% confidence is used as a metric. Hence, by selecting the s images (*S* ⊆ *U)* which the model is least confident about, the model performance is optimized^42^.

### Learning loss

Learning loss is a second network, called loss prediction module, which can be added to an active learning network, then called target model. It is trained to predict the losses of the target model on unlabeled inputs, simultaneously to the training of the target model. For the next active learning cycle this module can be used to select images for which the target model is likely to produce a wrong prediction^13^.

### BADGE

BADGE^10^ is an active learning method, which selects diverse samples that have a high magnitude in the gradient space. The model is considered to be uncertain about an image, if knowing the label of the image results in a large gradient of the loss with respect to model parameters. As the labels are not known, BADGE considers the predicted labels as true labels. Secondly, in order to make sure that a diverse batch of images are selected, BADGE uses the k-MEANS++ algorithm^43^. Hence, BADGE trades off between uncertainty and diversity of the s images *S* ⊆ *U* which are selected for active learning.

### Margin confidence sampling

Margin sampling is similar to least confidence sampling. Only for margin confidence sampling the difference between the most confident prediction and second most confident prediction is used as the metric. The main idea is that the smaller the difference is, the higher the model uncertainty on an image. As a result, the s images *S* ⊆ *U* with the least difference are selected^44^.

### Random sampling

During each active learning cycle an image set *S* ⊆ *U* is chosen arbitrarily. Random sampling acts as a baseline. Hence, all other algorithms are expected to perform better than random sampling.

### Pre-training methods

Network initialization can increase the performance of neural networks^45^. It is considered to be even more essential when the amount of annotated data is not considerably large^35^. In this work we utilize three different pre-training methods plus random initialization (baseline):

### ImageNet weights

ImageNet weights are obtained by training a feature extraction network on the ImageNet dataset. After training on ImageNet data, the weights of the feature extractor network can be used for initialization of models, which are to be trained on other datasets^46^. This has become a standard pre-training for classification tasks as it often helps the network to converge faster than with random initialization. Additionally, it has been shown to be beneficial in low-data biomedical imaging regimes^34^.

### Autoencoders

Autoencoders are a class of neural networks used for feature extraction^47^. The objective of the autoencoders is to reconstruct the input. An encoder network *e* encodes the input *x* into its latent representation *e*(*x*). The encoder typically includes a bottleneck layer with relatively few nodes. The bottleneck layer forces the encoder to represent the input data in a compact form. This latent representation is then used as an input to a decoder network *d,* which aims to output a reconstruction *d*(*e*(*x*)) of the original input. Hence, autoencoders do not require labels for training and the whole dataset can be used for training an autoencoder architecture. For pre-training the encoder is used as a feature extraction network while the decoder is generally discarded. This has been shown to significantly improve network initialization on biomedical imaging datasets^48^.

### SimCLR

SimCLR is a framework for contrastive learning of visual representations^14^. It learns representations in a self-supervised manner by using an objective function that minimizes the difference between representations of the model *f*^⊝^ on pairs of differently augmented copies of the same image. Let *a* be a function that performs stochastic data augmentations (such as cropping, adding color jitter, horizontal flipping and gray scale) on a given image. Each image *x* ∈ *D* in a mini-batch of size *B* is passed through the stochastic data augmentation function *a* twice to obtain *x*_*i*_ = {*x*_*1i*_’, *x*_*2i*_’}. These pairs can be termed as positive pairs as they originate from the same image *x*_*i*_. A neural network encoder *e* extracts the feature vectors *h* from the augmented images. A multi-layer perceptron with one hidden layer is used as a projection head for projecting the feature vectors *h* to the projection space where then, a contrastive loss is applied. The contrastive loss function is a softmax loss function applied on a similarity measure between positive pairs against all the negative examples in the batch and is weighted by the temperature parameter τ that controls the weight of negative examples in the objective function. Using SimCLR as a pre-training method shows significant improvement on ImageNet classification^14^.

### Random initialization

It has been shown that complete random initialization performs poorly compared to more sophisticated initialization measures^49^. We thus use Kaiming He initialization^50^ (which has been shown to boost the performance) as a baseline random initialization method.

### Training strategies

Large amounts of unlabeled data are typically available in biomedical applications. Ideally, this unlabeled data is not only used for network initialization but also during training. Thus, we compare the performance of training the model only using the existing labeled data a.k.a. supervised learning versus a semi-supervised approach, which incorporates the unlabeled data in the training process.

### Semi-supervised learning

For semi-supervised learning we use FixMatch^21^, a combination of consistency regularization^51^ and pseudo-labeling^52^. Given the set of unlabeled images *U* = {*u*_1_, *u*_2_, *u*_3_…*u*_*K*_} with |*U*| = *K*, consistency regularization aims to maximize the similarity between model outputs, obtained by passing stochastically augmented versions of the same image through the model *f*^⊝^(*a*(*x*)). Pseudo-labeling refers to using pseudo-labels for unlabeled images. Pseudo-labels are obtained by passing the unlabeled images through the model *f*^⊝^, i.e. *Y*^ = *f*^⊝^(*U*) and using the outcome with maximum probability in the predicted distribution *q*^_*i*_ = argmax *P*^⊝^(*y*^_*i*_|*x*_*i*_) as the pseudo-label if the maximum probability value *q*^_*i*_ is above a threshold *τ*. Using pseudo-labels, the unlabeled images are added to the set of labeled images *L* temporarily.

### FixMatch

The FixMatch loss consists of a supervised loss term i.e. the multi-class cross-entropy loss and the unsupervised loss term. The unsupervised loss term is calculated by passing the unlabeled dataset through a stochastic weak augmentation function *a*_weak_ (e.g. rotation or translation) and then applying pseudo-labeling on the output prediction distribution with threshold. Another set of pseudo-labels is obtained by passing the unlabeled dataset through a strong stochastic augmentation function *a*_strong_ (e.g. color distortion, random noise, or random erasing). After calculating the two sets of pseudo-labels for unlabeled images, consistency regularization is applied by calculating cross-entropy between the pseudo-labels. The loss function contains the weighting parameter λ which weighs the unsupervised loss term:

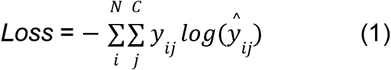

Using FixMatch, a significant performance improvement has been observed compared to supervised training in a low-data regime ^21^.

### Pseudo-labeling

The second method we use for semi-supervised learning is pseudo-labeling^53^. Wrapper methods involve training a base learner on *L* = {(*x*_*1*_, *y*_*1*_), (*x*_*2*_, *y*_*2*_), (*x*_*3*_, *y*_*3*_)…(*x*_*N*_, *y*_*N*_)} with |*L*| = *N* as well as *U* = {*u*_*1*_, *u*_*2*_, *u*_*3*_…*u*_*K*_} with |*U*| = K, for which the labels are acquired through pseudo-labeling^54^. The training process involves two steps. First, the base learner is trained on *L* as well as the pseudo-labeled set from previous cycles and predictions (y^). Second, the unlabeled images, for which the base learner outputs predictions with a high confidence, are assigned the corresponding predicted label and added to the training set as pseudo-labeled images for the next cycle.

### Supervised learning

In supervised learning we are looking for a model *f*^*⊝*^ with parameters ⊝ to learn a mapping *Y*^ = *f*^*⊝*^(*L*) such that the objective function *Loss* (*y*^_*i*_, *y*_*i*_) is minimized. Supervised learning uses only labeled data. The performance of the model can be evaluated using an evaluation metric *M* such as accuracy, recall etc. The objective function used in this paper is the multi-class cross-entropy loss function,

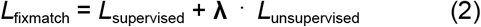

with *C* being the total number of classes in the dataset and *N* being the size of *L*.

### Architecture

We use ResNet18^55^ as the training architecture. For each dataset, we pretrain the ResNet18 using an autoencoder or SimCLR^14^. For the autoencoder pre-training, we use a feature extractor network consisting of a ResNet18 encoder and a decoder with transposed convolutional layers. After training the autoencoder, the ResNet18 encoder is used as a feature extractor and the decoder is discarded.

## Acknowledgements

We thank Björn Menze, Tingying Peng, Christian Matek, Melanie Schulz, Rudolf Matthias Hehr, Lea Schuh, Valerio Lupperger, and Ario Sadafi (Munich) for discussions and for contributing their ideas.

## Author contributions

ABQ implemented code and conducted experiments with supervision of SSB and DW. SSB, ABQ, DW, and CM wrote the manuscript with FS. SSB created figures with ABQ and the main storyline with CM. FS helped with the manuscript narrative and editing. CM supervised the study. All authors have read and approved the manuscript.

## Additional Information

### Data and Software availability

All scripts and how to access and process the data can be found here: https://github.com/marrlab/Med-AL-SSL.

